# Phenotypic profiling of macrocyclic lactones on parasitic *Schistosoma* flatworms

**DOI:** 10.1101/2022.09.12.507717

**Authors:** Kaetlyn T. Ryan, Nicolas J. Wheeler, Isaac K. Kamara, Hailey Johnson, Judith E Humphries, Mostafa Zamanian, John D. Chan

**Affiliations:** Department of Pathobiological Sciences, University of Wisconsin - Madison, Madison, WI, USA; Department of Chemistry, University of Wisconsin - Oshkosh, Oshkosh, WI, USA; Department of Biology, Lawrence University, Appleton, WI, USA

**Author notes:** Address correspondence to John D. Chan.

## Abstract

Macrocyclic lactones are front-line therapies for parasitic roundworm infections, but there are no comprehensive studies on the activity of this drug class against parasitic flatworms. Ivermectin is well known to be inactive against flatworms. However, the structure-activity relationship of macrocyclic lactones may vary across phyla, and it is entirely possible other members of this drug class do in fact show antiparasitic activity on flatworms. For example, there are several reports hinting at the anti-schistosomal activity of doramectin and moxidectin. To explore this class further, we developed an automated imaging assay combined with measurement of lactate levels from worm media. This assay was applied to the screening of 21 macrocyclic lactones (avermectins, milbemycins and others such as spinosyns) against adult schistosomes. These *in vitro* assays identified several macrocyclic lactones (emamectin, milbemycin oxime, and the moxidectin metabolite 23-ketonemadectin) that caused contractile paralysis and lack of lactate production. Several of these were also active against miracidia, a juvenile life cycle stage of the parasite. Hits prioritized from these *in vitro* assays were administered to mice harboring patent schistosome infections. However, no reduction in worm burden was observed. Nevertheless, these data show the utility of a multiplexed *in vitro* screening platform to quantitatively assess drug action and prioritize hits in a chemical series for *in vivo* studies. While the prototypical macrocyclic lactone ivermectin displays minimal activity against adult *Schistosoma mansoni*, this family of compounds does contain schistocidal compounds which may serve as a starting point for development of new anti-flatworm chemotherapies.

## Introduction

There is a recognized need for new anthelmintics given the limited treatment options for many parasitic infections and the widespread emergence of veterinary drug resistance (1). This problem is particularly acute for the neglected tropical disease schistosomiasis. The current frontline schistosomiasis monotherapy, praziquantel, has been in use for four decades. While praziquantel is typically between 70-90% effective (2), there are instances of treatment failure (3, 4). Parasite resistance has been selected in laboratory settings, and this is conceivably a risk in the field with more widespread administration of praziquantel monotherapy (5, 6). One barrier that contributes to the scarcity of leads in the development pipeline is that *in vitro* phenotypic screens on parasitic worms are often low throughput. The search for new anti-schistosomal compounds has often used juvenile schistosomula (7). However, juvenile schistosomes have fewer cells and few cell types (8–10), and they have differing sensitivity to some drugs (11). Adult parasites are the disease causing life cycle stage dwelling within the host, and scalable, quantitative methodologies for screening small molecules against these worms are needed to advance anti-schistosomal drug development.

In this study, we applied automated assays measuring the activity of a specific chemical series, macrocyclic lactones (ML), on adult schistosomes. MLs have a broad spectrum of activity, targeting ligand-gated ion channels (LGICs) of parasites across several phyla (roundworms and arthropods, reviewed in (12)), as well as numerous vertebrate LGICs (13–16). However, little is known about the structure - activity relationship of MLs in flatworms.

The most commonly used ML anthelmintic, ivermectin, is inactive against intramammalian parasitic flatworms (17–20). Ivermectin is not an agonist on schistosome GluCl isoforms (21). However, ivermectin B_1b_ does kill aquatic life cycle stages of schistosomes (22), and ivermectin does phenocopy praziquantel in several assays on free-living flatworms (23–25). Therefore, despite the lack of activity of ivermectin against intramammalian stages of parasites, this class of compounds is not inactive. Since the structure - activity relationships of MLs can vary depending on the parasite being studied (26), it is possible that other MLs do exhibit potent effects on schistosomes even if ivermectin is relatively inactive. For example, data from human clinical trials (27), screens against juvenile schistosomula (28) and computational predictions of antischistosomal drugs (29) have indicated moxidectin may have anti-schistosomal effects. These data indicate that at least some MLs do exhibit activity against flatworms, even if they do not act through the GluCl channels that MLs typically target. Therefore, we sought to comprehensively assess the structure - activity relationship of MLs on adult schistosomes using a panel of several phenotypic assays.

## Results

### In vitro measurement of drug activity on adult schistosomes

*In vitro* phenotypic screening of worms can be challenging because there is not a single outcome that is a reliable predictor of impaired viability within the host *in vivo*. Worm movement is often used as an initial screening outcome, reasoning that dead worms will not exhibit movement. Past methods of quantitatively analyzing adult schistosome movement were manual and low throughput, but methods using optical flow algorithms have been developed for automated screening of parasite movement (30–32). These approaches allow for quantitative readouts not biased by subjective scoring, but on their own do not necessarily provide a readout of worm viability since paralyzed worms may still be alive and recover after drug treatment. Assays benefit from a secondary readout of drug activity, such as lactate measurements from conditioned media which provides a readout of metabolic activity (6, 33). Therefore, we sought to combine these approaches in a multiplexed assay for parasite viability (Figure 1A). We first optimized assay conditions using two drugs known to paralyze worms *in vitro*, meclonazepam and dichlorvos, and then proceeded to screen 21 commercially available MLs.

**Figure 1.**
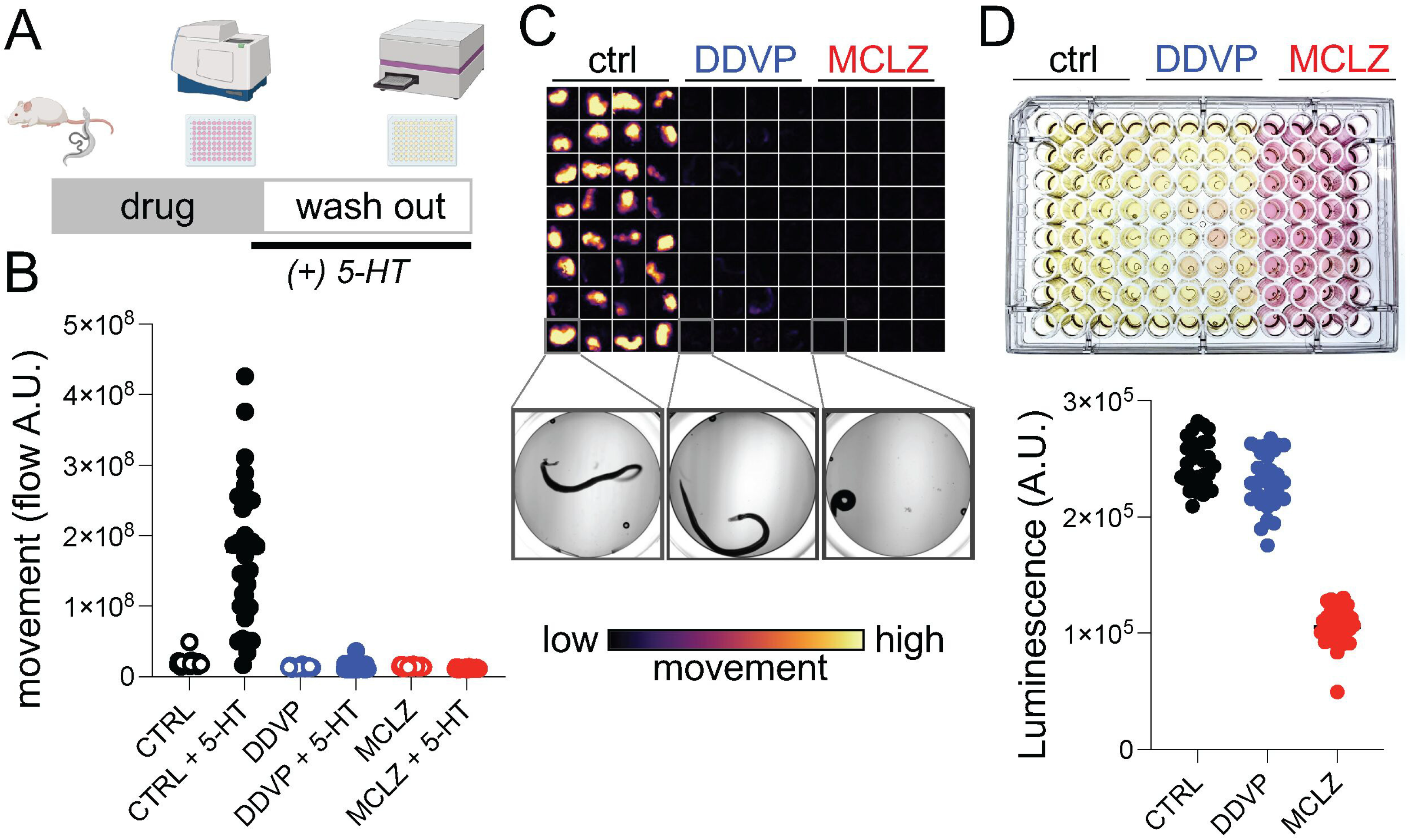
Multiplexed assay for schistosome viability. **(A)** Drug screening workflow on adult *S. mansoni*. Worms are harvested from mice, exposed to test compound for 24 hours (3 columns or 32 worms / drug), supplemented with 5-HT and imaged to record movement (15 sec / well). Media was changed to fresh drug-free media for follow up assay measuring lactate production. **(B)** Quantification of movement for worms exposed to either DMSO control, dichlorvos (DDVP, 10 µM), or meclonazepam (MCLZ, 10 µM). Worms were imaged before (open symbols) or after (closed symbols) serotonin addition (5-HT, 250 µM). **(C)** Flow cloud showing worm movement over the period of recording for the plate. Yellow areas indicate worm movement, while dark wells remain paralyzed. Below = brightfield images of an individual worm representative of each treatment. **(D)** *Top -* Plate media after culture (24hrs in 5-HT, 250 µM) to allow accumulation of lactate. *Bottom -* measurement of lactate from the same plate using the Lactate-Glo assay system.

Adult parasites were harvested from the mesenteric vasculature of mice 6-7 weeks post-infection and allowed to unpair at room temperature. Sexually mature schistosomes live as pairs with the female worm residing within the male ventral gynecophoral canal, although worms tend to unpair during *in vitro* culture. To obtain separate male and female populations for subsequent assays, unpaired female worms were removed from the petri dish and cultured separately. The rational for using unpaired worms is that if a male and female worm were to unpair in a well, the movement values may be roughly double that of wells with a single male worm. Additionally, female worms lay many eggs (up to ∼100 per day) which will accumulate in the well. These eggs will move as worms thrash in the media, producing high motility values since the flow algorithms will detect any pixel that is moving, regardless of whether it is worm or debris.

After harvesting, worms were treated with meclonazepam (MCLZ, 10 µM), which is known to produce contractile paralysis (34), and the cholinergic compound dichlorvos (DDVP, 10 µM), an acetylcholinesterase inhibitor known to produce flaccid paralysis (35, 36). From visual observation, it was clear that untreated control worms were motile, but the movement was too slow to record within a recording window that allowed for convenient scaling across multiple wells in a plate. Within the recording period of 15 seconds per well, only a roughly 20% difference in control and drug treated worms was detectable. Therefore, to expand the signal window between DMSO control and drug treated worms, we supplemented the wells with serotonin (5-HT, 250 μM) for one hour prior to recording worm movement (Figure 1B). Treatment with 5-HT increased the baseline motility of worms >8 fold relative to control worms without 5-HT. Under these conditions, DDVP and MCLZ inhibited movement by 92% and 93% respectively, relative to 5-HT treated controls (Figure 1B&C).

After motility recordings (Figure 1B), drug containing media was removed and replaced with fresh 5-HT containing media (200 µL per well). Lactate production can be visually observed by acidification of media and yellowing of phenol red pH indicator, and levels were quantified using the Lactate-Glo luminescence assay (Figure 1D). Worms treated with MCLZ appeared killed by drug treatment and lactate measurements were 58% less than those of control wells. DDVP treated worms appear to be merely paralyzed but not killed, since these worms produced lactate at levels comparable to control worms (only a 7% decrease, Figure 1D). The differences between these two compounds, which both inhibit worm movement, illustrate the utility of using a workflow with multiple readouts to assess drug activity.

### In vitro screen of macrocyclic lactones

Using this multiplexed, quantitative assay of worm movement and metabolic activity, we proceeded to profile the activity of commercially available MLs against *S. mansoni*. These compounds included various avermectins (which contain a carbohydrate attached to the macrocyclic lactone ring system) and milbemycins (which lack the carbohydrate), as well as several other members of the class that are more structurally divergent (Figure 2A, Supplemental Table 1). Adult parasites were incubated in various concentrations of compound (1 - 20 μM) for 24 hours. While most worms did not exhibit changes in morphology with ML treatment, several treatments caused a coiled, contractile phenotype (emamectin, doramectin, milbemycin oxime, milbemycin D, 23-ketonemadectin, spinosad; Figure 2B and Supplemental Figure 1). Worms were then imaged in 96 well plates in media supplemented with 5-HT (250 μM), using the motility assay described above. Concentration-response data are summarized in Figure 2A, with black symbols denoting the concentration at which worm motility was decreased by over two thirds relative to untreated controls. Compounds that caused obvious changes to worm morphology generally also were effective at inhibiting worm movement relative to controls. However, several compounds that did not cause clear contractile phenotypes were also nonetheless effective at inhibiting movement. For example, eprinomectin treated worms appeared morphologically normal but their movement was inhibited at drug concentrations 5 µM and above. Full concentration-response curves for all compounds tested are shown in Supplemental Figure 2.

**Figure 2.**
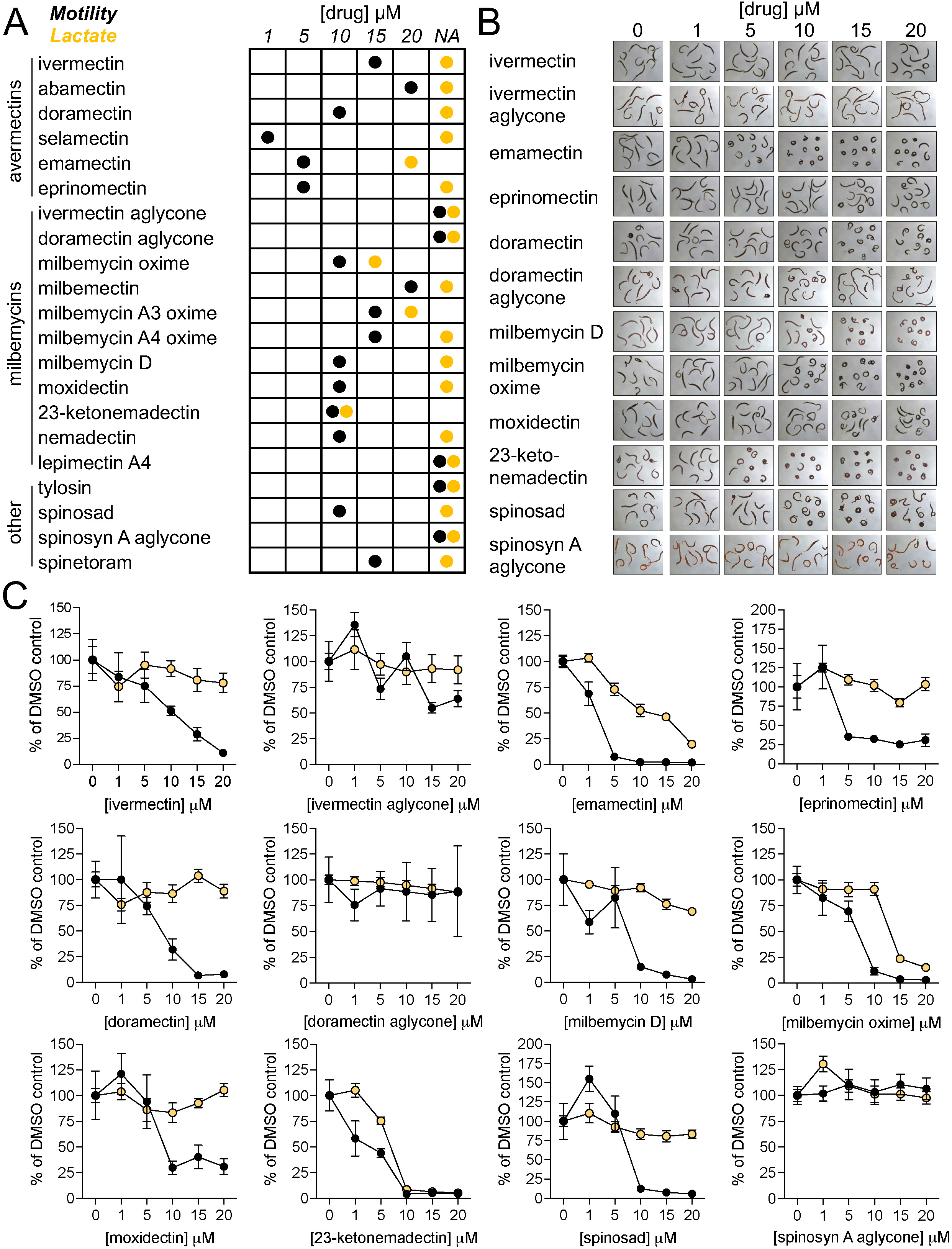
*In vitro* effect of macrocyclic lactones on adult S. mansoni. **(A)** Summary of concentration-response screening data from motility and lactate assays across 21 different macrocyclic lactones. Adult male *S. mansoni* were cultured in drug for 24 hours (1 column or 8 worms per drug concentration) and then assayed for movement and lactate production as per Figure 1. Motility is shown in black symbols and lactate assays in orange, reflecting the threshold where movement is less than 30% of control. **(B)** Images of adult male worms treated with various concentrations of drug, showing concentration-dependent effects on worm morphology for certain compounds. **(C)** Concentration-response curves for compounds shown in B, plotted with motility (black) and lactate (orange) reflecting mean ± standard error. Concentration-response curves for all 21 compounds listed in (A) can be found in the Supplemental Figure **2**.

Following imaging of worm movement, drug containing media was replaced with fresh, drug-free culture media supplemented with 5-HT (250 μM). Plates were incubated for a further 24-48 hours, and then media was harvested to measure lactate production (Figure 2A&C). Lactate concentration-response curves were right-shifted relative to the motility readout, indicating that worms are capable of a degree of recovery after drug-washout. Treatments that caused contractile paralysis typically also inhibited lactate production at concentrations that corresponded to maximal movement inhibition. For example, 23-ketonemadectin caused complete contractile paralysis of worms at 10 μM and also completely inhibited movement and lactate production at this concentration. However, at lower concentrations movement was partially inhibited while lactate remained relatively unchanged. Finally, not all treatments that caused contractile paralysis killed worms. For example, spinosad and doramectin treated worms displayed clear changes in morphology (Figure 2B) and movement was inhibited (Figure 2C), but following drug washout these worms recovered and produced lactate at levels comparable to controls at all concentrations tested.

In other organisms, MLs act through activation of inhibitory LGICs (37). Another compound that works through activation of inhibitory LGICs and has activity on *S. mansoni* is the benzodiazepine meclonazepam (38). However, meclonazepam does not exhibit activity against all species of schistosomes - *Schistosoma japonicum* are unaffected by this compound (34). Therefore, we were interested in whether MLs may also be inactive against this species. Adult *S. japonicum* were harvested from mice 7 weeks post-infection and treated with a selection of compounds that displayed activity on *S. mansoni* (23-ketonemadectin, emamectin, milbemycin oxime and doramectin). All hits that were active against *S. mansoni* also displayed activity against *S. japonicum*, causing contractile paralysis (Figure 3A) and inhibition of motility and lactate production (Figure 3B). These data indicate that MLs may be capable of targeting multiple species of parasites.

**Figure 3.**
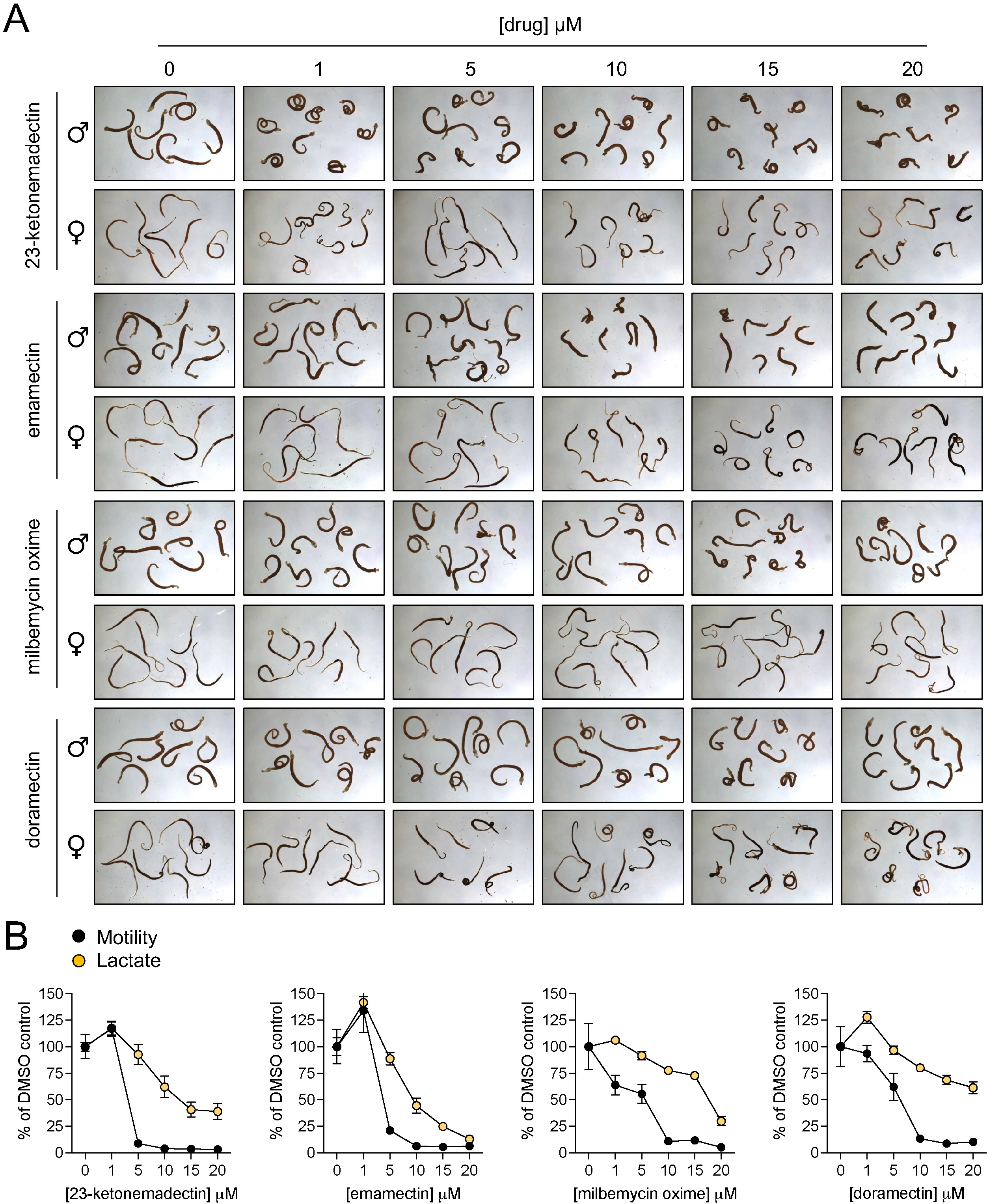
Activity of macrocyclic lactones on adult *S. japonicum*. **(A)** Morphology of adult male and female worms treated with selected macrocyclic lactones active on *S. mansoni* (23-ketonemadectin, emamectin, milbemycin oxime, doramectin). **(B)** Concentration - response curves for motility (black) and lactate production (orange) of worms treated with drugs shown in (A). Symbols reflect mean ± standard error.

The activity of MLs on adult schistosomes is of clear interest because this life cycle stage dwells within infected humans. However, one of the few published reports of ML activity on schistosomes is on juvenile stages and the snail intermediate host (22). To determine whether the activity profile we observed in adults may also hold for other life-cycle stages, we performed a motility assay on *S. mansoni* miracidia hatched from eggs isolated from the livers of *S. mansoni* infected mice. Miracidia were aliquoted into 24 well plates and incubated in drug-containing artificial pond water. A selection of MLs were screened at a fixed concentration of 5 μM. This concentration was chosen because it represents an intermediate concentration at which more potent compounds displayed activity against adult worms (ex. 23-ketonemadectin) while less active compounds showed no discernible effect (ex. ivermectin). Videos of worm motility (30 seconds each) were recorded after a one hour drug incubation period. Praziquantel (3 μM) was chosen as a positive control since this concentration was previously reported to kill miracidia (39). The image stacks from the acquired videos were processed to provide a single minimum intensity projection for each well, providing a proxy for worm movement over the recording period. Paths of individual miracidia can be inferred from dark trails across the images (Figure 4). This provides a qualitative readout of whether worms in the wells exhibit movement or not. Because miracidia also swim rapidly up and down in the Z axis, entering and leaving the focal plane, quantitative analysis of individual worm movement from these videos is more difficult. However, these minimum intensity projections allow for simple scoring of whether miracidial movement does or does not occur for a given treatment. The results were consistent across replicates that milbemycin oxime and 23-ketonemadectin were active against miracidia (0/6 replicates moving for each). Ivermectin and doramectin, which were less effective against adult worms, did not obviously reduce movement relative to DMSO controls.

**Figure 4.**
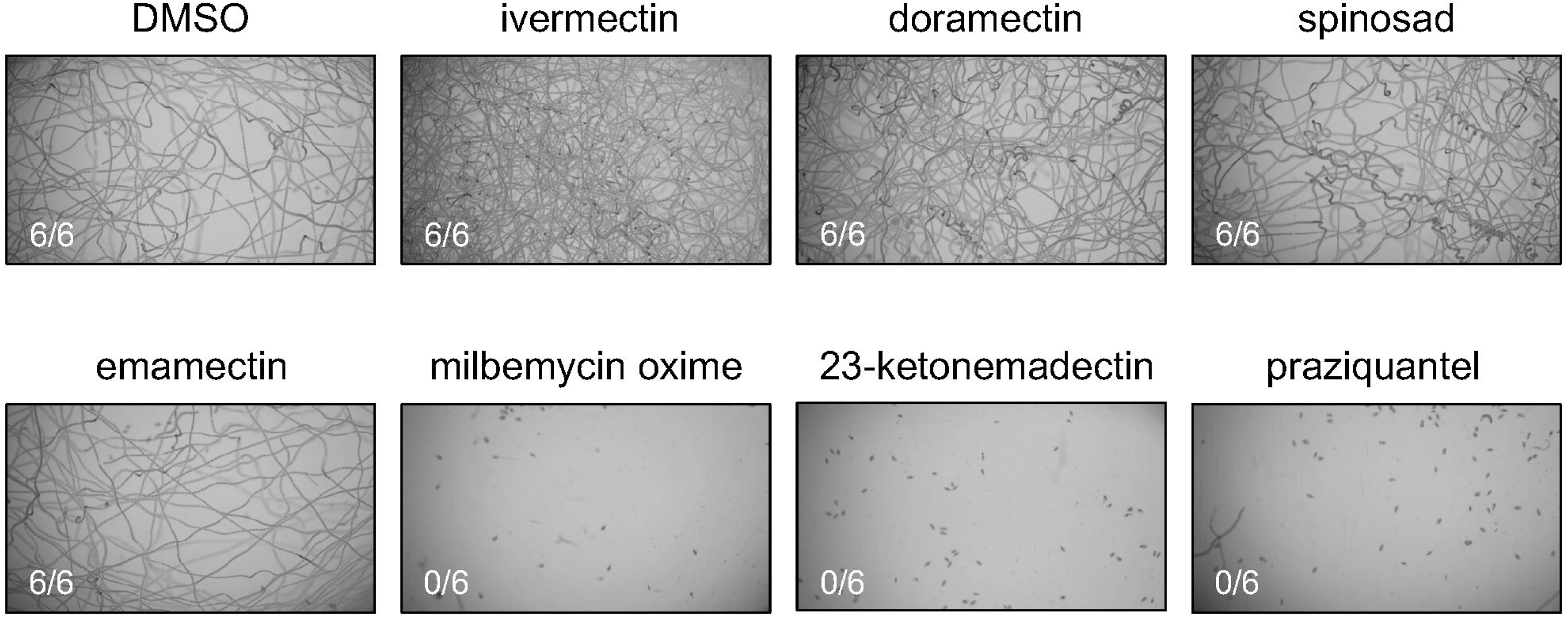
Activity of macrocyclic lactones against *S. mansoni* miracidia. Freshly hatched *S. mansoni* miracidia were incubated in either DMSO, test compound (5 μM of each ML), or praziquantel (3 μM) for one hour. Movement was recorded for 30 seconds, and the resulting image stacks were processed to produce the maximum intensity projections shown above. Each drug treatment was repeated as 6 technical replicates, the scored number shown reflects wells with worm movement.

### In vivo activity of macrocyclic lactones against S. mansoni

We proceeded to test the *in vivo* efficacy of several MLs in a murine model of schistosomiasis. Prior studies on the ML doramectin reported that this drug reduced *S. mansoni* parasite burden in mice by 60% (28). In addition to doramectin, we chose to test emamectin, 23-ketonemadectin and milbemycin oxime because of the activity of these compounds in motility and lactate assays. Ivermectin was selected as a negative control as it has been extensively reported as inactive against schistosomes, and eprinomectin was also selected due to the structural similarity to emamectin. Female Swiss Webster mice harboring six week *S. mansoni* infections were administered compounds by oral gavage. Doses were chosen to be as close to the maximum tolerated dose as possible while not exhibiting toxic effects. Animals were sacrificed one week later to count parasite burden. No reduction in worm burden was observed for any treatment (Figure 5), despite the relatively high doses of compound administered.

**Figure 5.**
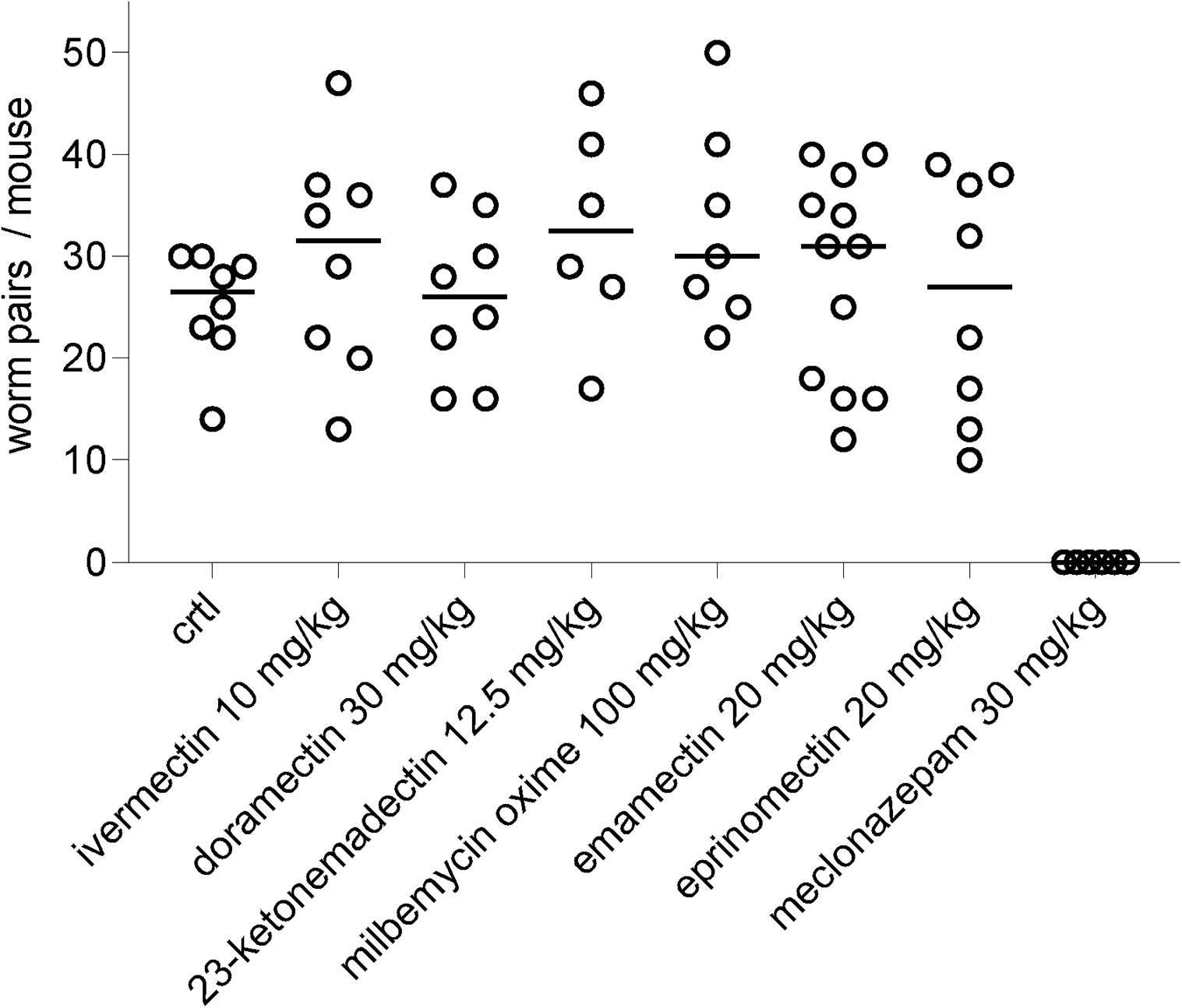
*In vivo* efficacy of macrocyclic lactones against adult *S. mansoni*. Mice harboring patent schistosome infections were treated orally with either vehicle negative control, various MLs, or meclonazepam (positive control) at the dose indicated. Open symbol = worm burden of an individual mouse, solid bar = mean worm burden.

## Discussion

Macrocyclic lactones (MLs) are considered ineffective against parasitic flatworm infection, in large part due to studies that have shown ivermectin to be inactive on these worms (17–19). However, there are scattered reports of other MLs that do exhibit activity on flatworms *in vitro* or *in vivo* (22, 23, 27, 28). To explore this chemical series further, we screened 21 MLs (avermectins, milbemycins, spinosyns) and identified several MLs active *in vitro* against *S. mansoni* and *S. japonicum*.

### Quantitative multiplexed assay for movement and metabolic activity

We looked at several different endpoints to assess ML activity, and the profile of different compounds varied across these assays. Motility assays tended to be more sensitive, 16/21 compounds inhibited movement to some degree at the concentrations tested (Figure 2A). These include many treatments which did not exhibit obvious changes in worm morphology such as curling/contraction. However, inhibited movement does not necessarily indicate a particularly deleterious effect of drug on the parasite. Following drug washout, many worms seem quite capable of recovering. This is indicated by the right shift in potency of the concentration - response curves in the lactate measurement assay relative to the motility assays (Figure 2, Supplemental Figure 2). In fact, some treatments that caused near complete inhibition of movement had no effect on lactate production after washout (ex. eprinomectin, doramectin, spinosad). A few compounds did show activity in all endpoints - emamectin, milbemycin oxime and 23-ketonemadectin caused contractile paralysis and inhibition of movement and lactate production. Several MLs have been screened against schistosomes as part of a larger library (28), but the activity of these three compounds has not been previously reported.

Notably, all of the compounds tested were inactive *in vivo* (Figure 5). This included doramectin, which displayed subcurative efficacy in another murine model (28). However, the Liberian strain of *S. mansoni* and NMRI line of mice used in that study differ from the parasite and host strains provided by the NIH-NIAID Schistosomiasis Resource Center (NMRI *S. mansoni* in Swiss Webster mice). Variability in drug screening results between laboratories that use different parasite and mouse strains has been reported (28, 40). Our *in vitro* data (Figures 2 & 3) show that compounds exhibit activity in the low micromolar range when incubated for 24 hours. However, it is likely that the free-drug concentration *in vivo* that worms are exposed to does not reach this level for this duration of time. This issue of whether *in vitro* drug concentrations can be therapeutically achieved *in vivo* has also been raised regarding the repurposing of ivermectin as a SARS-CoV-2 therapy. Ivermectin exhibits anti-viral effects *in vitro* at 5 µM, but *in vivo* only reaches sub-micromolar plasma concentrations - and even then, much of the drug is protein bound (41). Nevertheless, our data still show that MLs as a class can exhibit schistocidal activity. This study screened only 21 compounds against schistosomes, while hundreds of MLs have been screened to yield the nematicidal compounds currently in use (26). Further exploration of this chemical series is likely to produce more potent hits.

### Structure-activity relationship of MLs on S. mansoni

Consistent with prior studies, ivermectin did not exhibit schistocidal activity. However, other members of the class were capable of killing worms *in vitro* (Figure 2). These were not restricted to one group of compounds (i.e. just avermectins or just milbemycins), or molecules with specific functional. Generally, the structure-activity relationship reproduced several observations reported for MLs on other organisms. Some observations can be made regarding substituents on the carbohydrate, benzofuran and spiroketal rings.

Several aglycones, which are missing the carbohydrate ring system, were tested and all were inactive (Figure 2C). This is consistent with data from nematodes showing that the aglycones, which contain a polar hydroxy group on the C-13 position, are inactive while the milbemycins, which also lack the sugar molecule at this position, retain activity (26). Comparison of emamectin, eprinomectin and abamectin is also useful for determining the importance of substitution at the carbohydrate C4 position. Emamectin was more capable of killing worms than eprinomectin. Emamectin contains a methylamino group at the carbohydrate C4 position, while eprinomectin contains an acetylamino group. This is in contrast to abamectin, which contains a hydroxy group at this position and exhibits even less activity (Figure 2A, Supplemental Figure 2).

The activity of milbemycin oxime, and to a lesser extent selamectin, indicates schistosomes may be amenable to substitution of the C5 position of the benzofuran ring with an oxime. These compounds displayed comparable activity to others with a C5 hydroxy group. This is in contrast to nematodes, which are less tolerant of C5 substitutions (26, 42).

Modification of the spiroketal ring system also alters drug activity. Differences in schistocidal and molluscicidal activity have been reported for ivermectin; ivermectin B1_a_ contains a sec-butyl group and is relatively inactive (unlike what is observed in nematodes) while ivermectin B1_b_ contains a isopropyl group on the spiroketal ring and is more active (22). A particularly interesting pair of compounds from this study is moxidectin and its metabolite 23-ketonemadectin. Moxidectin did not cause contractile paralysis of worms. While it did partially inhibit movement, it had no impact on lactate production. Treatment with 23-ketonemadectin, which differs from moxidectin in that it possesses a ketone group rather than a methoxyamine at the C23 position of the spiroketal ring, caused contractile paralysis, inhibition of movement and lack of lactate production. Given the schistocidal activity of 23-ketonemadectin *in vitro*, it is possible that this metabolite contributes to the 70% *S. mansoni* egg reduction rate observed in humans treated with moxidectin (27).

### ML mechanism of action on flatworms

These *in vitro* data raise the question of what is the target and mechanism of action for MLs in flatworms that underpins paralysis and death? In nematodes, MLs target inhibitory LGICs while also requiring connections between adjacent cells / tissues to propagate the nematicidal effects (43). Even if LGICs are relatively conserved across roundworms and flatworms and MLs are also acting on inhibitory schistosome LGICs, flatworm and nematode anatomy is quite different and so the drug mechanisms are likely to differ between the two phyla.

MLs certainly have a wide range of targets - some are LGICs and some are not (44, 45). The flatworm target of these compounds is likely not a GluCl as it is in nematodes. The GluCl ivermectin binding pocket requires a glycine residue at a specific transmembrane domain (46). Schistosomes contain a phenylalanine or isoleucine at this position - both of which have large side chains that make this pocket inaccessible to drug (47). This has been confirmed by cloning and heterologous expression of *S. mansoni* GluCls, which are insensitive to ivermectin (21).

However, consideration of potential targets does not need to be restricted to GluCls, since MLs are relatively promiscuous ligands that have been shown to act on numerous different types of LGICs (13, 14, 16, 48, 49).

In addition to GluCls, schistosomes possess acetylcholine-gated chloride channels (ACCs)(50), nicotinic acetylcholine receptors (nAchRs) which are predicted to be cationic (51), and a clade of unique channels not found outside of Lophotrochozoa which lack the Cys-loop motif that typically defines LGICs (52). If MLs were to be acting on a schistosome LGIC, cationic nAchRs are potential targets to be explored. In insects, a nAchR is the target of the spinosyns (53–57), which in schistosomes cause the same contractile phenotype as the avermectins and milbemycins (Figure 2B). This phenotype is more consistent with cation influx than the action of inhibitory channels, which may be expected to cause flaccid paralysis. Studies on recombinantly expressed human α7nAChRs and α4β2 nAChRs found them to be sensitive to emamectin, but not ivermectin (58), which would be consistent with emamectin having greater activity on schistosomes (Figure 2). A separate ML binding site within nAchRs has been proposed, an ‘intra-subunit’ site, that is distinct from the inter-subunit site that is blocked by large schistosome amino acid side chains (47). The potency of avermectin-type MLs on nAchRs is also lower, in the micromolar range, which is more in line with the concentrations of drug used in this study than the concentrations typically active on GluCls. These data point to nAchRs as interesting targets to explore, although functional expression of parasite nAchRs is not trivial as this can involve iterating through heteromeric subunit combinations and may require various accessory subunits (59).

## Materials and methods

### Harvesting adult schistosomes

Female Swiss Webster mice infected with either *S. mansoni* (NMRI strain) or *S. japonicum* (Philippine strain) were sourced from the NIH-NIAID Schistosomiasis Resource Center. Mice were euthanized by CO_2_ asphyxiation between 6-7 weeks post-infection and adult schistosomes were harvested from the mesenteric vasculature. Livers were set aside in ice-cold 1.2 % NaCl and saved for miracidia hatching as mentioned below. Adult worms were placed in culture media consisting of high-glucose DMEM supplemented with 5% fetal calf serum, penicillin-streptomycin (100 units/mL), HEPES (25 mM) and sodium pyruvate (1 mM). Worms were treated in test compounds and cultured at 37°C / 5% CO_2_ for 24 hours prior to conducting motility assays. All animal work, including *in vivo* drug treatments outlined below, were carried out with the oversight and approval of UW-Madison Research Animal Resources and Compliance (RARC), adhering to the humane standards for the health and welfare of animals used for biomedical purposes defined by the Animal Welfare Act and the Health Research Extension Act. Experiments were approved by the UW-Madison School of Veterinary Medicine IACUC committee (approved protocol #V006353-A08).

### Sourcing test compounds

Compounds were sourced as follows: dichlorvos (MilliporeSigma), meclonazepam (Toronto Research Chemicals), ivermectin (Alfa Aesar), abamectin (Thermo Scientific), doramectin (Selleck Chemicals, Thermo Scientific), selamectin (Cayman Chemical, MedChemExpress), emamectin (Santa Cruz Biotechnology), eprinomectin (MilliporeSigma), ivermectin aglycone (Cayman Chemical), doramectin aglycone (Cayman Chemical), milbemycin oxime (MedChemExpress, TargetMol), milbemectin (Cayman Chemical), milbemycin A3 oxime (Cayman Chemical), milbemycin A4 oxime (Cayman Chemical), milbemycin D (Toronto Research Chemicals), moxidectin (TCI America), 23-ketonemadectin (Toronto Research Chemicals, United States Biological), nemadectin (Cayman Chemical), lepimectin A4 (Cayman Chemical), tylosin (Ambeed), spinosad (Cayman Chemical), spinosyn A aglycone (Cayman Chemical), spinetoram (Cayman Chemical). Structures for compounds screened are provided in Supplemental Table 1.

### Adult schistosome motility assays

Worms harvested from mice and incubated in either DMSO control or test compounds listed above were imaged in clear 96 well plates on an ImageXpress Nano (Molecular Devices). Except for experiments shown in Figure 1B that state otherwise, serotonin (5-HT) was added at 250 µM to increase worm movement at least 1 hour prior to imaging. Time lapse recordings were acquired for each well (15 second videos at a rate of 4 frames / second) using a 2X objective. Movement was analyzed using wrmXpress as outlined in (32). On occasion, individual wells can produce unusually high flow values due to debris or eggs (if paired worms have been inadvertently selected) that are moving in recording as well. Since worm movement should already be close to maximal with addition of 5-HT, high flow readings >3 standard deviations over the mean flow values for a given drug treatment were removed.

### Lactate assays

Following imaging on the ImageXpress Nano, media was then removed from each well using a multichannel pipette and replaced with fresh culture media supplemented with 5-HT (250 µM). Depending on the volume of media placed in the well, the time for lactate accumulation varied. The optimal period was 1 day culture in 200 µL media per well, since prolonged culturing increases the likelihood of microbial contamination. Media was removed and stored in sealed plates at -80°C, until lactate was measured with the Lactate-Glo assay kit (Promega) using solid white half area 96-well plates. In order to test within the dynamic range of the assay kit, all samples were diluted 1:250 in PBS.

### Miracidia motility assays

Livers harvested from schistosome-infected mice were processed to retrieve eggs and hatch miracidia as described previously (60). Ten livers were processed at a time. Briefly, livers were washed in saline (1.2% NaCl), homogenized using a stainless-steel blender for 1 minute. The volume of the resulting homogenate was brought to approximately 800 mL with fresh saline and centrifuged at 290 x g for 15 min at 4°C. The pellet was resuspended in saline and the wash step repeated, and the resulting purified eggs were resuspended in artificial pond water (final salt concentrations of CaCO_3_ 66.6 µM, MgCO_3_ 7.9 µM, NaCl 11.4 µM, KCl 1.8 µM) in a 1-L volumetric flask covered with foil up to the neck and hatching was facilitated with a light source directed towards the flask neck. Concentrated hatched miracidia were collected from the top layer of pond water, and the volume was brought to 24 mL with pond water. Miracidia were then aliquoted 0.5mL per well in 24 well plates (approximately 300 miracidia per well) and treated with test compounds for 1 hour. Videos (30 seconds) were recorded using a stereomicroscope (Zeiss Stemi 508 with Axiocam 208 camera), converted to avi format using FFmpeg and processed in ImageJ.

### In vivo drug screening

At 6 weeks post-infection, *S. mansoni* infected mice were administered a single dose of test compound (ivermectin 10 mg/kg, doramectin 30 mg/kg, 23-ketonemadectin 12.5 mg/kg, milbemycin oxime 100 mg/kg, emamectin 20 mg/kg, eprinomectin 20 mg/kg, meclonazepam 30 mg/kg) solubilized in vegetable oil (0.25 mL) and delivered by oral gavage. These doses were selected based on the toxicity of each compound. Mice were euthanized as outlined above at 7 weeks post-infection and worms were dissected from the liver and mesenteries to determine parasite burden for each mouse.

## Acknowledgments

Schistosome infected mice were provided by the NIH-NIAID Schistosomiasis Resource Center for distribution through BEI Resources, NIH-NIAID Contract HHSN272201700014I. This work was supported by funding from NIH-NIAID R21AI146540 (JDC), R21AI153545 (JDC & MZ) and 5T32GM081061-04 (KTR).

---

**Supplemental Figure 1. Morphology of schistosomes treated with macrocyclic lactones**. Morphology of adult male and female *S. mansoni* treated with 21 different macrocyclic lactones for 24 hours.

**Supplemental Figure 2. Motility and lactate production of schistosomes treated with macrocyclic lactones**. Motility and lactate production of adult male *S. mansoni* treated with 21 different macrocyclic lactones for 24 hours. Concentration - response curves plotted with motility (black) and lactate (orange), reflecting mean ± standard error.

**Supplemental Table 1. Structures of macrocyclic lactones screened in this study**. Names and structures (SMILES IDs) for all compounds screened.

## Works Cited

1. Nixon SA, Welz C, Woods DJ, Costa-Junior L, Zamanian M, Martin RJ. 2020. Where are all the anthelmintics? Challenges and opportunities on the path to new anthelmintics. Int J Parasitol Drugs Drug Resist 14:8–16.

2. Ismail M, Metwally A, Farghaly A, Bruce J, Tao LF, Bennett JL. 1996. Characterization of isolates of Schistosoma mansoni from Egyptian villagers that tolerate high doses of praziquantel. Am J Trop Med Hyg 55:214–218.

3. Danso-Appiah A, De Vlas SJ. 2002. Interpreting low praziquantel cure rates of Schistosoma mansoni infections in Senegal. Trends Parasitol 18:125–129.

4. Kron M, Gordon C, Bauers T, Lu Z, Mahatme S, Shah J, Saeian K, McManus DP. 2019. Persistence of Schistosoma japonicum DNA in a Kidney-Liver Transplant Recipient. Am J Trop Med Hyg 100:584–587.

5. Fallon PG, Doenhoff MJ. 1994. Drug-resistant schistosomiasis: resistance to praziquantel and oxamniquine induced in Schistosoma mansoni in mice is drug specific. Am J Trop Med Hyg 51:83–88.

6. Le Clec’h W, Chevalier FD, Mattos ACA, Strickland A, Diaz R, McDew-White M, Rohr CM, Kinung’hi S, Allan F, Webster BL, Webster JP, Emery AM, Rollinson D, Djirmay AG, Al Mashikhi KM, Al Yafae S, Idris MA, Moné H, Mouahid G, LoVerde P, Marchant JS, Anderson TJC. 2021. Genetic analysis of praziquantel response in schistosome parasites implicates a transient receptor potential channel. Sci Transl Med 13:eabj9114.

7. Chen S, Suzuki BM, Dohrmann J, Singh R, Arkin MR, Caffrey CR. 2020. A multi-dimensional, time-lapse, high content screening platform applied to schistosomiasis drug discovery. Commun Biol 3:747.

8. Diaz Soria CL, Lee J, Chong T, Coghlan A, Tracey A, Young MD, Andrews T, Hall C, Ng BL, Rawlinson K, Doyle SR, Leonard S, Lu Z, Bennett HM, Rinaldi G, Newmark PA, Berriman M. 2020. Single-cell atlas of the first intra-mammalian developmental stage of the human parasite Schistosoma mansoni. Nat Commun 11:6411.

9. Wendt G, Zhao L, Chen R, Liu C, O’Donoghue AJ, Caffrey CR, Reese ML, Collins JJ 3rd. 2020. A single-cell RNA-seq atlas of Schistosoma mansoni identifies a key regulator of blood feeding. Science 369:1644–1649.

10. Wangwiwatsin A, Protasio AV, Wilson S, Owusu C, Holroyd NE, Sanders MJ, Keane J, Doenhoff MJ, Rinaldi G, Berriman M. 2020. Transcriptome of the parasitic flatworm Schistosoma mansoni during intra-mammalian development. PLoS Negl Trop Dis 14:e0007743.

11. Reimers N, Homann A, Hoschler B, Langhans K, Wilson RA, Pierrot C, Khalife J, Grevelding CG, Chalmers IW, Yazdanbakhsh M, Hoffmann KF, Hokke CH, Haas H, Schramm G. 2015. Drug-induced exposure of Schistosoma mansoni antigens SmCD59a and SmKK7. PLoS Negl Trop Dis 9:e0003593.

12. Martin RJ, Robertson AP, Choudhary S. 2021. Ivermectin: An Anthelmintic, an Insecticide, and Much More. Trends Parasitol 37:48–64.

13. Sigel E, Baur R. 1987. Effect of avermectin B1a on chick neuronal gamma-aminobutyrate receptor channels expressed in Xenopus oocytes. Mol Pharmacol 32:749–752.

14. Krause RM, Buisson B, Bertrand S, Corringer P-J, Galzi J-L, Changeux J-P, Bertrand D. 1998. Ivermectin: A Positive Allosteric Effector of the α7 Neuronal Nicotinic Acetylcholine Receptor. Mol Pharmacol 53:283–294.

15. Khakh BS, Proctor WR, Dunwiddie TV, Labarca C, Lester HA. 1999. Allosteric Control of Gating and Kinetics at P2X4Receptor Channels. J Neurosci 19:7289–7299.

16. Shan Q, Haddrill JL, Lynch JW. 2001. Ivermectin, an unconventional agonist of the glycine receptor chloride channel. J Biol Chem 276:12556–12564.

17. Campbell WC, Fisher MH, Stapley EO, Albers-Schönberg G, Jacob TA. 1983. Ivermectin: a potent new antiparasitic agent. Science 221:823–828.

18. Campbell WC. 1991. Ivermectin as an antiparasitic agent for use in humans. Annu Rev Microbiol 45:445–474.

19. Shoop WL, Ostlind DA, Rohrer SP, Mickle G, Haines HW, Michael BF, Mrozik H, Fisher MH. 1995. Avermectins and milbemycins against Fasciola hepatica: in vivo drug efficacy and in vitro receptor binding. Int J Parasitol 25:923–927.

20. Vicente B, López-Abán J, Chaccour J, Hernández-Goenaga J, Nicolas P, Fernández-Soto P, Muro A, Chaccour C. 2021. The effect of ivermectin alone and in combination with cobicistat or elacridar in experimental Schistosoma mansoni infection in mice. Sci Rep 11:4476.

21. Dufour V, Beech RN, Wever C, Dent JA, Geary TG. 2013. Molecular cloning and characterization of novel glutamate-gated chloride channel subunits from Schistosoma mansoni. PLoS Pathog 9:e1003586.

22. Katz N, Araújo N, Coelho PMZ, Morel CM, Linde-Arias AR, Yamada T, Horimatsu Y, Suzuki K, Sunazuka T, Ōmura S. 2017. Ivermectin efficacy against Biomphalaria, intermediate host snail vectors of Schistosomiasis. J Antibiot 70:680–684.

23. Nogi T, Zhang D, Chan JD, Marchant JS. 2009. A Novel Biological Activity of Praziquantel Requiring Voltage-Operated Ca2+ Channel β Subunits: Subversion of Flatworm Regenerative Polarity. PLoS Negl Trop Dis 3:e464.

24. Simanov D, Mellaart-Straver I, Sormacheva I, Berezikov E. 2012. The Flatworm Macrostomum lignano Is a Powerful Model Organism for Ion Channel and Stem Cell Research. Stem Cells Int 2012:167265.

25. Ferenc NN, Levin M. 2019. Effects of Ivermectin Exposure on Regeneration of D. dorotocephala Planaria: Exploiting Human-Approved Ion Channel Drugs as Morphoceuticals. Macromol Biosci 19:e1800237.

26. Shoop WL, Mrozik H, Fisher MH. 1995. Structure and activity of avermectins and milbemycins in animal health. Vet Parasitol 59:139–156.

27. Barda B, Coulibaly JT, Puchkov M, Huwyler J, Hattendorf J, Keiser J. 2016. Efficacy and Safety of Moxidectin, Synriam, Synriam-Praziquantel versus Praziquantel against Schistosoma haematobium and S. mansoni Infections: A Randomized, Exploratory Phase 2 Trial. PLoS Negl Trop Dis 10:e0005008.

28. Panic G, Vargas M, Scandale I, Keiser J. 2015. Activity Profile of an FDA-Approved Compound Library against Schistosoma mansoni. PLoS Negl Trop Dis 9:e0003962.

29. Zorn KM, Sun S, McConnon CL, Ma K, Chen EK, Foil DH, Lane TR, Liu LJ, El-Sakkary N, Skinner DE, Ekins S, Caffrey CR. 2021. A Machine Learning Strategy for Drug Discovery Identifies Anti-Schistosomal Small Molecules. ACS Infect Dis 7:406–420.

30. Marcellino C, Gut J, Lim KC, Singh R, McKerrow J, Sakanari J. 2012. WormAssay: a novel computer application for whole-plate motion-based screening of macroscopic parasites. PLoS Negl Trop Dis 6:e1494.

31. Wheeler NJ, Heimark ZW, Airs PM, Mann A, Bartholomay LC, Zamanian M. 2020. Genetic and functional diversification of chemosensory pathway receptors in mosquito-borne filarial nematodes. PLoS Biol 18:e3000723.

32. Wheeler NJ, Gallo KJ, Garncarz EJ, Ryan KT, Chan JD, Zamanian M. 2022. wrmXpress: A modular package for high-throughput image analysis of parasitic and free-living worms. bioRxiv.

33. Howe S, Zophel D, Subbaraman H, Unger C, Held J, Engleitner T, Hoffmann WH, Kreidenweiss A. 2015. Lactate as a novel quantitative measure of viability in Schistosoma mansoni drug sensitivity assays. Antimicrob Agents Chemother 59:1193–1199.

34. Stohler HR. 1978. Ro 11-3128, a novel schistosomicidal compound. Proceedings of the 10th International Congress of Chemotherapy 1:147–148.

35. Barker LR, Bueding E, Timms AR. 1966. The possible role of acetylcholine in Schistosoma mansoni. Br J Pharmacol Chemother 26:656–665.

36. Mellin TN, Busch RD, Wang CC, Kath G. 1983. Neuropharmacology of the parasitic trematode, Schistosoma mansoni. Am J Trop Med Hyg 32:83–93.

37. Wolstenholme AJ, Neveu C. 2022. The avermectin/milbemycin receptors of parasitic nematodes. Pestic Biochem Physiol 181:105010.

38. McCusker P, Mian MY, Li G, Olp MD, Tiruveedhula VVNPB, Rashid F, Golani LK, Verma RS, Smith BC, Cook JM, Chan JD. 2019. Non-sedating benzodiazepines cause paralysis and tissue damage in the parasitic blood fluke Schistosoma mansoni. PLoS Negl Trop Dis 13:e0007826.

39. Andrews P. 1978. Effect of praziquantel on the free living stages of Schistosoma mansoni. Z Parasitenkd 56:99–106.

40. Maccesi M, Aguiar PHN, Pasche V, Padilla M, Suzuki BM, Montefusco S, Abagyan R, Keiser J, Mourão MM, Caffrey CR. 2019. Multi-center screening of the Pathogen Box collection for schistosomiasis drug discovery. Parasit Vectors 12:493.

41. Peña-Silva R, Duffull SB, Steer AC, Jaramillo-Rincon SX, Gwee A, Zhu X. 2021. Pharmacokinetic considerations on the repurposing of ivermectin for treatment of COVID-19. Br J Clin Pharmacol 87:1589–1590.

42. Michael B, Meinke PT, Shoop W. 2001. Comparison of ivermectin, doramectin, selamectin, and eleven intermediates in a nematode larval development assay. J Parasitol 87:692–696.

43. Dent JA, Smith MM, Vassilatis DK, Avery L. 2000. The genetics of ivermectin resistance in Caenorhabditis elegans. Proc Natl Acad Sci U S A 97:2674–2679.

44. Laing R, Gillan V, Devaney E. 2017. Ivermectin - old drug, new tricks? Trends Parasitol 33:463–472.

45. Chen I-S, Kubo Y. 2018. Ivermectin and its target molecules: shared and unique modulation mechanisms of ion channels and receptors by ivermectin. J Physiol 596:1833–1845.

46. Lynagh T, Lynch JW. 2010. A glycine residue essential for high ivermectin sensitivity in Cys-loop ion channel receptors. Int J Parasitol 40:1477–1481.

47. Lynagh T, Lynch JW. 2012. Ivermectin binding sites in human and invertebrate Cys-loop receptors. Trends Pharmacol Sci 33:432–441.

48. Collins T, Millar NS. 2010. Nicotinic acetylcholine receptor transmembrane mutations convert ivermectin from a positive to a negative allosteric modulator. Mol Pharmacol 78:198–204.

49. Zheng Y, Hirschberg B, Yuan J, Wang AP, Hunt DC, Ludmerer SW, Schmatz DM, Cully DF. 2002. Identification of two novel Drosophila melanogaster histamine-gated chloride channel subunits expressed in the eye. J Biol Chem 277:2000–2005.

50. MacDonald K, Buxton S, Kimber MJ, Day TA, Robertson AP, Ribeiro P. 2014. Functional characterization of a novel family of acetylcholine-gated chloride channels in Schistosoma mansoni. PLoS Pathog 10:e1004181.

51. Bentley GN, Jones AK, Oliveros Parra WG, Agnew A. 2004. ShAR1α and ShAR1β: novel putative nicotinic acetylcholine receptor subunits from the platyhelminth blood fluke Schistosoma. Gene 329:27–38.

52. Jaiteh M, Taly A, Hénin J. 2016. Evolution of Pentameric Ligand-Gated Ion Channels: Pro-Loop Receptors. PLoS One 11:e0151934.

53. Perry T, McKenzie JA, Batterham P. 2007. A Dalpha6 knockout strain of Drosophila melanogaster confers a high level of resistance to spinosad. Insect Biochem Mol Biol 37:184–188.

54. Puinean AM, Lansdell SJ, Collins T, Bielza P, Millar NS. 2013. A nicotinic acetylcholine receptor transmembrane point mutation (G275E) associated with resistance to spinosad in Frankliniella occidentalis. J Neurochem 124:590–601.

55. Somers J, Nguyen J, Lumb C, Batterham P, Perry T. 2015. In vivo functional analysis of the Drosophila melanogaster nicotinic acetylcholine receptor Dα6 using the insecticide spinosad. Insect Biochem Mol Biol 64:116–127.

56. Zimmer CT, Garrood WT, Puinean AM, Eckel-Zimmer M, Williamson MS, Davies TGE, Bass C. 2016. A CRISPR/Cas9 mediated point mutation in the alpha 6 subunit of the nicotinic acetylcholine receptor confers resistance to spinosad in Drosophila melanogaster. Insect Biochem Mol Biol 73:62–69.

57. Lu W, Liu Z, Fan X, Zhang X, Qiao X, Huang J. 2022. Nicotinic acetylcholine receptor modulator insecticides act on diverse receptor subtypes with distinct subunit compositions. PLoS Genet 18:e1009920.

58. Xu X, Sepich C, Lukas RJ, Zhu G, Chang Y. 2016. Emamectin is a non-selective allosteric activator of nicotinic acetylcholine receptors and GABAA/C receptors. Biochem Biophys Res Commun 473:795–800.

59. Noonan JD, Beech RN. 2022. Reconstitution of an N-AChR from Brugia malayi. bioRxiv.

60. Dinguirard N, Heinemann C, Yoshino TP. 2018. Mass Isolation and In Vitro Cultivation of Intramolluscan Stages of the Human Blood Fluke Schistosoma Mansoni. J Vis Exp https://doi.org/10.3791/56345.

